# Decoding continuous variables from event-related potential (ERP) data with linear support vector regression (SVR) using the Decision Decoding Toolbox (DDTBOX)

**DOI:** 10.1101/2021.05.31.446502

**Authors:** Stefan Bode, Elektra Schubert, Hinze Hogendoorn, Daniel Feuerriegel

## Abstract

**Background:** Multivariate classification analysis for event-related potential (ERP) data is a powerful tool for predicting cognitive variables. However, classification is often restricted to categorical variables and under-utilises continuous data, such as response times, response force, or subjective ratings. An alternative approach is Support Vector Regression (SVR), which uses single-trial data to predict continuous variables of interest.

**New Method:** In this tutorial-style paper, we demonstrate how SVR is implemented in the Decision Decoding Toolbox (DDTBOX). To illustrate in more detail how results depend on specific toolbox settings and data features, we report results from two simulation studies resembling real EEG data, and one real ERP-data set, in which we predicted continuous variables across a range of analysis parameters.

**Results:** Across all studies, we demonstrate that SVR is effective for analysis windows ranging from 2 ms – 100 ms, and relatively unaffected by temporal averaging. Prediction is still successful when only a small number of channels encode true information, and the analysis is robust to temporal jittering of the relevant information in the signal.

**Comparison with existing Methods:** Our result show that SVR as implemented in DDTBOX can reliably predict continuous, more nuanced variables, which may not be well-captured by classification analysis.

**Conclusions:** In sum, we demonstrate that linear SVR is a powerful tool for the investigation of single-trial EEG data in relation to continuous variables, and we provide practical guidance for users.

## Introduction

### 1.1 General Background

Multivariate analysis techniques for non-invasively acquired neuroimaging data with a high temporal resolution, such as electroencephalography (EEG) and magnetoencephalography (MEG) data, have become increasingly popular in cognitive neuroscience research. The use of classifiers by means of multivariate pattern analysis (MVPA) in particular has the advantage that it can potentially extract more information from the signal at a given point in time than would be possible using classical univariate approaches. This can lead to insights about what information or cognitive process is represented in brain activity patterns at very short timescales and can shed light on the dynamic development of neural representations over time (Carlson et al., 2020; Contini et al., 2017; King & Dehaene, 2014).

In recent years, several toolboxes have been published that allow researchers to apply these techniques to EEG/MEG data, including ADAM (Fahrenfort et al., 2018), CoSMoMVPA (Oosterhof et al., 2016), DDTBOX (Bode et al., 2019), FieldTrip (Oostenveld et al., 2011), MNE-Python (Gramfort et al., 2013), MVPA-Light (Treder, 2020), and the neural decoding toolbox (Meyers, 2013), complementing similar toolboxes for functional magnetic resonance imaging (fMRI; e.g., Hanke et al., 2009; Hebart et al., 2015; Schrouff et al., 2013). In addition, several tutorial-style papers have been published that provide practical advice for users (e.g., Bode et al., 2019; Carlson et al., 2020; Fahrenfort et al., 2018; Grootswagers et al., 2016; Treder, 2020).

An important constraint of multivariate pattern classifiers, as they are most frequently implemented, is that they are restricted to predicting categorical variables, because they use multivariate signals to assign trials to distinct classes. Recently, there has been a growing interest in the prediction of continuous variables from multivariate M/EEG signals for which classification analyses are therefore not well-suited. For example, researchers have attempted to discriminate between high and low values of a continuous variable using median splits (e.g., Korjus et al., 2015), however this approach often suffers from reduced statistical power (additional issues are discussed in MacCallum et al., 2002).

An alternative approach is Support Vector Regression (SVR), which can be used to predict continuous variables of interest from single-trial data, such as response times, response force, subjective ratings (e.g., emotional state, valence, etc.), and any other variable that can be associated with EEG patterns (e.g., Bode et al., 2014; Lan et al., 2016; Schubert et al., 2020; Siswandari et al., 2019). This approach is particularly useful if one is interested in decoding behaviourally meaningful continuous variables that cannot easily (or without loss of nuanced information) be divided into distinct classes (e.g., Sabbagh et al., 2020; for applications using Magnetoencephalography (MEG) data see e.g., Trübutschek et al., 2017; and for applications with EEG data beyond the prediction of cognitive variables see, e.g., Li et al., 2008; Jach et al., 2020; Sato et al., 2008).

In this article, we will describe the implementation of SVR in the Decision Decoding Toolbox (DDTBOX), which has increasingly been used to decode different cognitive processes, ranging from low-level vision to higher level abstract representations, from EEG amplitude data (Bode et al., 2019). The aim of this paper is to first provide potential users with background information about what types of questions have been (and can be) addressed using SVR followed by practical guidance on how to implement such analyses in the toolbox. We note that there are other implementations of this and similar approaches (e.g., Li et al., 2008; Lan et al., 2016; Sato et al., 2008). Note that our paper is not intended as a comprehensive comparison between different multivariate approaches (e.g., linear discriminant analysis, random forest classification), nor to demonstrate superiority of one approach over another. Instead, we focus on one specific implementation in DDTBOX here, which is designed to provide users with code that is easy to adapt to new research questions. We further note that there are many aspects and variants of SVR (and parameters in DDTBOX) which will also not be covered (e.g., the use of non-linear kernels), and users will be referred to our extensive online Wiki. The chosen parameters and features of data, however, map onto to the most frequently encountered analytical decisions in cognitive neuroscience research, and we believe that users will benefit most from the overview and the analyses we provide here.

We first explain the general principles of conducting SVR in DDTBOX, including a brief overview of some analysis parameters, which can be adjusted by the user. These include a) the features included in the analysis, i.e. whether a *spatial* or a *spatiotemporal* analysis is conducted; and b) the choice of an *appropriate window width* for the analysis window that is moved through the trial to capture potential information in the signal. This section serves to provide users with a guide on how to best utilise the toolbox. We then briefly review the types of cognitive processes that have been investigated using SVR. Note that to maximise comparability with the approach, rather than providing a comprehensive review of all multivariate regression studies, we again focus on studies that have used DDTBOX.

In the subsequent section, we present an analysis of simulated EEG data to showcase the changes in results if the key parameters are varied. For this, we created a dataset for 37 simulated participants (this *N* corresponded to our own recent EEG study, which is also analysed here) with 100 trials each (simulating noisy amplitude data, sampled at 500 Hz), time-locked to an event-of-interest (as is common for ERP analyses). In each trial, we injected signal into random activity for a circumscribed number of channels during a small time-window (as it typically occurs in ERP studies investigating brief cognitive processes). The signal scaled with a simulated continuous cognitive variable in each trial (which could be e.g., rating scale values, response times, or any other variable that can meaningfully be measured at interval levels). We subjected this data to the SVR function of DDTBOX and used all combinations of spatial and spatiotemporal analyses with separate analyses using analysis time windows of 2 ms, 10 ms, 20 ms, 50 ms, and 100 ms. We show that across all combinations, the information could be reliably decoded. The results further revealed that there was no strong advantage for using the (computationally more demanding) *spatiotemporal* analysis over the simpler *spatial* analysis (which neglects temporal information within the analysis time window). Furthermore, it was found that some temporal smearing (i.e. a forward projection) of information occurred with larger time windows (50 ms and 100 ms width).

In the third section, we present the results of a second simulation study in which we focussed on spatial SVR using the same analysis time window widths as before. This time, we systematically varied a) the *temporal jitter* (i.e. the variance) with which the signal was present in the data, and b) the *number of channels* that contained any signal. This served to showcase the impact of two variations of signal that can be expected from real data on the decodability of the information. These analyses showed that more temporal variability in information and less informative channels led to the expected drop in decodability; however, information could still be recovered from the data in each case.

Notably, the simulation studies constituted an idealised, simplistic case of information representation: there was always signal in each trial, and the signal always scaled with the variable-of-interest. Additionally, the way that the variable was ‘represented’ in the signal was simplified as it was present in all signal-carrying channels alike, which does not take into account more complex multivariate coding schemes that may be present in real signal patterns. In the final section, we therefore demonstrate that SVR can also recover information from real data for which we do not know exactly how information is represented. For this, we reanalysed data from a recent study (Schubert et al., 2021) that was comparable in size to the simulated data. In this study, in each trial participants rated the tastiness and healthiness of a visually presented food item. We again used spatial and spatiotemporal SVR in combination with all analysis window widths to decode both ratings in separate analyses, demonstrating highly similar results to the simulated data.

Finally, we summarise the findings, discuss limitations of these studies, and end with recommendations for users on how to tailor SVR in DDTBOX for their purposes, and a short discussion on the use of SVR in general.

### 1.2 Support Vector Regression (SVR) analysis in DDTBOX

#### 1.2.1 Implementation of SVR analyses in DDTBOX

The latest version of DDTBOX (Version 1.0.5) allows users to perform either support-vector machine (SVM) classification, interfacing with LIBSVM (Chang & Lin, 2011) or LIBLINEAR (Fan et al., 2008), or support vector regression (SVR; interfacing with LIBSVM) to analyse EEG amplitude data (note that it can also be used to analyse other formats, such as spectral power data; e.g., Jach et al., 2020; but this option is not yet routinely integrated). The epsilon-insensitive linear SVR method as implemented by default in DDTBOX confers many of the advantages of SVMs to perform regression based on multivariate patterns of EEG data. In contrast to standard linear regression, in epsilon SVR any residuals (errors) of less than a set value of epsilon are ignored, and only residuals larger than this value determine the structure of the regression model (for further details see Hastie et al., 2009).

Before running the SVR analysis, the data is pre-processed in the same way as for a classical ERP analysis. It has been suggested that data cleaning can be less stringent for MVPA given that, for example, noisy and non-informative channels and unsystematic artefacts would not hurt the classifier, because low weights will be assigned to those features during the classification (Carlson et al., 2020; Grootswagers et al., 2016); however, we prefer applying the same strict artefact rejection procedures to the data as for ERP analyses. This will also allow for making the data fully comparable to results from classical ERP analyses, which are often reported alongside the MVPA results. Users may potentially choose to perform current source density (CSD) analyses as a final pre-processing step. This method will not be reviewed in detail here (and is not performed for the reported data). In short, for CSD analysis, a Laplacian filter is applied to re-reference the data to the surrounding electrodes. This has the advantages that the data becomes independent from a specific reference channel, and the unique contributions from each channel are amplified while redundancies in the data are attenuated (Pernier, Perrin, & Bertrand, 1988; Perrin, Bertrand, & Pernier, 1987). The use of similar Laplacian filters has been suggested to improve classification (Bai et al., 2007). The resulting higher topographical accuracy of the CSD signals (Gevins, 1989), due to the reduction of redundancies in the signals from adjacent electrode sites, could indeed be beneficial also for pattern classification analysis using SVR (Bode et al., 2012).

For the SVR, the pre-processed data (time-locked to an event of interest) is then exported into a MATLAB data matrix with the format: channels x data points x trials. A second matrix, in the form of a single column containing the variable of interest for each trial (corresponding to the trials included in the EEG data matrix), is also generated. Each participant’s matrices serve as the input for a within-participant SVR to predict the variable-of-interest from distributed patterns of EEG amplitude data.

DDTBOX uses a moving-window approach in which the trial data (usually containing a baseline period and epoched and truncated, depending on the individual research question) is analysed within an *analysis time window*, which is moved through the entire trial in small (overlapping or non-overlapping) steps, each time containing the next step’s data. It is also possible to use a pre-defined time-period of interest instead (e.g., Billing et al., 2018), but we will focus on the moving-window approach in this paper. Each analysis step/window is treated as an independent analysis. In DDTBOX, a cross-validation procedure is applied for which the trials are randomly divided into different sets (e.g., ten sets for a ten-fold cross-validation). All sets but one are used for training, and the independent left-out data set is used for testing how well the trained regression model generalises to unseen data (for further details see Bode et al., 2019). This procedure is repeated for each fold of the cross-validation by using each data set once for testing while independently training on all other sets. In addition, DDTBOX allows for implementing multiple iterations of the entire cross-validation procedure, each time with a fresh, random re-sorting of trials into new sets (the default in DDTBOX is ten of such iterations of a ten-fold cross-validation). This step increases the overall time and computational processing costs, but it substantially decreases the probability of any false positive results that might by chance result from the initial sorting of data. The SVR outputs a Fisher-Z transformed correlation coefficient for the correlation between the real “labels” (i.e. the value of the variable-of-interest in each trial) and the predicted “label” (i.e. the predicted value of the variable-of-interest). The average result of all iterations of all cross-validation steps is the final output and assigned to the respective analysis time window. An identical analysis is then repeated for the data from each analysis time window until the end of the trial (i.e. the last analysis window) is reached (Figure 1).

**Figure 1:**
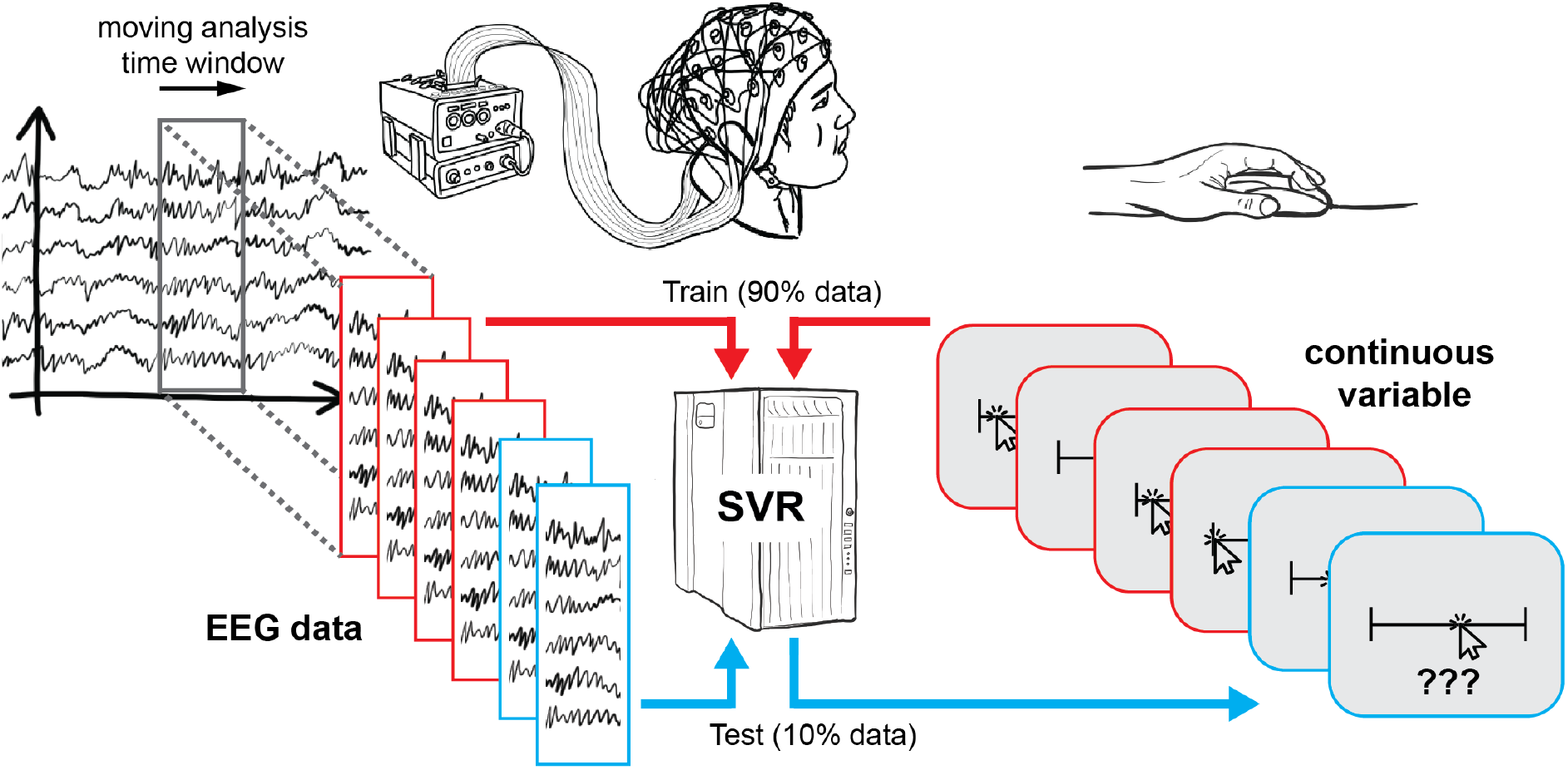
Schematic overview of Support Vector Regression. At each timestep, EEG data and continuous responses from 90% of trials are used to train a Support Vector Regression, which is then tested using the remaining 10% of trials. This procedure is then repeated such that each subset of trials is tested once.

Note that to perform the SVR, DDTBOX interacts with LIBSVM (Chang & Lin, 2011). It possible to choose different kernels for the SVR, but we will focus on the default setting, which is used in a great majority of cognitive neuroscience research of this kind, the linear kernel. LIBSVM further allows the user to change various other settings, such as the epsilon and cost parameters, which will not be considered here in detail.

Participants’ individual results are then submitted to group-level statistical testing. The default option in DDTBOX is to test the results of each analysis time window independently against an empirical chance distribution for the same time window, obtained by repeating an identical number of iterations of the same cross-validation procedure for each participant, with exactly the same data and the same labels, the only difference being that the assignment of labels to data is randomised (and again freshly randomised for each iteration of the cross-validation procedure). This constitutes a more conservative approach than testing against theoretical chance level (Combrisson & Jerbi, 2015), and it allows for controlling for any biases inherent in the data (for details see Bode et al., 2019). Decoding results from each analysis time window can then be tested for statistical significance using either paired-samples t-tests or a group-level analysis method (Allefeld et al., 2016) based on the minimum statistic (Friston et al. 1999). Several options for corrections for multiple comparisons can be used, some of which exploit the temporal autocorrelation of the classification results across analysis time windows to preserve statistical power, e.g., cluster-based permutation tests (Bullmore et al., 1999; Maris & Oostenveld 2007).

#### 1.2.2 Specific parameter settings for SVR in DDTBOX

To initially set up the SVR analysis, DDTBOX first requires the user to modify a MATLAB script that defines all input parameters for the specific data set (including the location of data on the computer, the number of channels, the sampling rate, etc.) as well as the parameters for the analysis to be performed.

The first decoding analysis parameter requires users to choose between a *spatial, temporal,* or *spatiotemporal* SVR analysis. We will neglect the temporal analysis here as it does not make use of the full spatial pattern of signal (see Bode et al., 2019 for details). The difference between the spatiotemporal and spatial analysis is that the spatiotemporal method extracts all available data points *within* the analysis time window for all (or all specified) channels as features (e.g., if data was recorded at 500 Hz, and the analysis time window width is 10 ms, and the data set contains 64 channels, this results in: 5 x 64 = 320 features). The spatial analysis, on the other hand, first averages across the data points for each channel *within* this analysis time window, resulting in only one feature per channel that represents the average signal for each channel for the respective window (e.g., 1 x 64 = 64 features for the same 10 ms analysis time window).

Next, the user is required to specify the *width of the analysis time window* and the step size with which the analysis time window is moved through the trial. If both are the same, then the analysis time window is moved in non-overlapping steps, while if the step size is smaller than the window width, overlapping analysis time windows are used that sample from partly the same data. This, in turn, has to be considered for the interpretation; but for the sake of this paper, we will simply use a step size of 10 ms. The most commonly used analysis time window widths range from 1 data point (2 ms for 500 Hz, or 1 ms for 1,000 Hz) to 100 ms windows, and we will systematically explore these window widths here.

There are multiple other settings the user can change, including whether the data should be normalised before decoding (the default setting is to normalise the data; normalising can also substantially reduce computation time), and whether feature weights should be extracted. We will not cover these here but refer to the toolbox Wiki for more details (https://github.com/DDTBOX/DDTBOX/wiki). The group-level analysis script also allows choosing different options for statistical tests, as mentioned above, but we will solely focus on standard group-level statistical tests using a series of paired-samples t-tests, corrected for multiple comparisons using cluster-based permutation tests based on the cluster mass statistic.

### 1.3 Research examples for SVR analyses using DDTBOX

Several published studies have successfully used SVR to decode behaviourally meaningful continuous variables from EEG data. For example, Bode and colleagues (2014) have used SVR in DDTBOX to predict post-experiment ratings of semantic attributes for rewarding objects during passive viewing. They recorded 64-channel EEG at 512 Hz and used a linear spatiotemporal SVR with 40 ms analysis time windows for prediction. The study found that it was possible to predict, from activity patterns recorded during the first 200 ms after passive exposure to these images, the post-experimental arousal ratings as well as ratings for how strongly the particular image was related to the participants’ individual concepts of the present or the future. Another possibility is to predict continuous aspects of behaviour, for example aspects of response force. Siswandari and colleagues (2019) asked their participants to produce a specific force pulse in each trial to match a given target range. Instead of only decoding whether the target range would be hit or not, they used linear spatiotemporal SVR to predict the specific force parameters (i.e. the peak force and time-to-peak) from EEG data recorded in the lead-up to response execution. They recorded from 61 channels measured at 500 Hz and used 10 ms analysis time windows for the analyses. Their results demonstrate that the neural signals during the early preparation period already contained specific information about the subsequently unfolding force response, beyond a simple error-signal. In another study by Schubert and colleagues (2020), linear SVR was used to predict the success ratings for different emotion regulation strategies from ERP signals extracted from the anticipation phase and the implementation phase of an emotion regulation task. They used 64-channel ERPs, recorded at 512 Hz, and applied a 20 ms analysis time window for prediction. The results showed that the degree to which participants were able to successfully apply a reappraisal strategy to modulate emotion intensity was predictable from early stages of the anticipation phase (i.e. when participants could anticipate the strategy use but were not yet shown a negative emotional stimulus), while the degree of distraction success was predictable from early stages of the implementation phase (i.e. when participants could actually apply the distraction strategy to regulate their negative emotional response). Finally, another study by Schubert and colleagues (2021) reported results from two experiments in which linear SVR was used to predict ratings of the healthiness and tastiness of visually presented snack food items. In the first experiment (from which the data were re-analysed for the present paper), participants gave the ratings immediately after being exposed to the items. In the second experiment, these ratings were given before the experiment, and EEG data was analysed from the subsequent phase of making consumption decisions. They recorded from 64 channels, recorded at 512 Hz, and used 20 ms time windows for prediction. In both studies, it was possible to predict healthiness and tastiness ratings from the distributed patterns of activity.

Taken together, these examples provide evidence that SVR is a powerful tool for the prediction of (even abstract) aspects of processed stimuli, detailed assessments of one’s own mental states, as well as fine-grained aspects of unfolding motoric responses. It is important to note that there is no one-size-fits-all approach, because different cognitive processes will require slightly different parameters to be uncovered, and hence the cited studies have used different parameters for the reported analyses. Some cognitive processes of interest might occur early and are brief, while others occur later and are sustained. These processes will, in turn, have a variety of different neural drivers, and might be reflected by very different ERPs in space and time. In the following sections, we will experimentally explore some of these aspects and show how different parameter settings, as well as variations in the signal, may impact the SVR results.

## 2. Simulation Study 1: SVR analysis type and analysis window width

### 2.1 Methods

#### 2.1.1 Data

All code and data used in the Simulation studies will be made available (at https://osf.io/ef4an/) at the time of publication. For each condition, we simulated 37 datasets (matching the sample size of the study by Schubert et al. (2021) that was reanalysed using identical procedures and is reported below), consisting of 100 epochs (corresponding to a rather typical-to-large number of trials in real experiments) spanning -100 ms to 1000 ms relative to event (i.e. simulated stimulus) onset. The number of channels was 64, and the sampling rate was 500 Hz. To generate noise in the EEG signal, for each channel and each trial we summed together multiple sinusoids with periodicities ranging between 1-40 Hz (in steps of 0.1 Hz) with randomised phases. The amplitude of each sinusoid was scaled so that higher frequencies were of smaller amplitude (multiplied by [1 / the sinusoid frequency in Hz]). The first 100 ms of the epoch were treated as a pre-stimulus baseline, and the resulting epochs were baseline-corrected using the average amplitude of this 100 ms baseline interval. This approach was used to impose a degree of temporal autocorrelation as found in real EEG data; however, similar results could also be obtained by generating Gaussian noise independently at each time point.

In addition to the noise, we systematically added “signal” to 8 channels during a specific time period. The signal was generated by adding a Gaussian-shaped curve to the noise for each of the 8 specified channels. The peak (i.e., time point of maximum amplitude) of the Gaussian was located at 400 ms from stimulus onset. The Gaussian standard deviation was 20 ms, meaning that 95% of the added signal was located within ±40 ms of the peak time point. The peak amplitude of the signal (i.e. the height of the Gaussian) scaled linearly with the value of the continuous variable that comprised the SVR condition label in each trial. We further varied the peak time points of the Gaussian-shaped signals across trials according to a boxcar distribution, in order to emulate temporal variability in EEG signals. We chose a jitter of ±30 ms, meaning that the peak of the signal Gaussian in a given trial was equally likely to occur between 370-430 ms. Continuous values comprising the SVR condition labels were generated by taking random draws from a Gaussian distribution with a mean of 0 and a standard deviation of 1. We used various settings of the SVR analysis in DDTBOX to predict the continuous variable from the multivariate data as described below.

#### 2.2.2 SVR analysis

Linear SVR in DDTBOX (Version 1.0.5) was used, interfacing LIBSVM (using default settings: epsilon-insensitive SVR algorithm; cost parameter C = 0.1). We analysed data separately applying A) spatial SVR, and B) spatiotemporal SVR. Within each of these analysis streams, we ran all analyses separately with different analysis window width: 2 ms, 10 ms, 20 ms, 50 ms, and 100 ms. The 2 ms analysis time window equates to one single data-point. As this is the smallest possible analysis window width, it naturally cannot contain temporal information within the analysis window (and was therefore counted as a spatial analysis only). Table 1 shows the different analysis conditions.

**Table 1:**
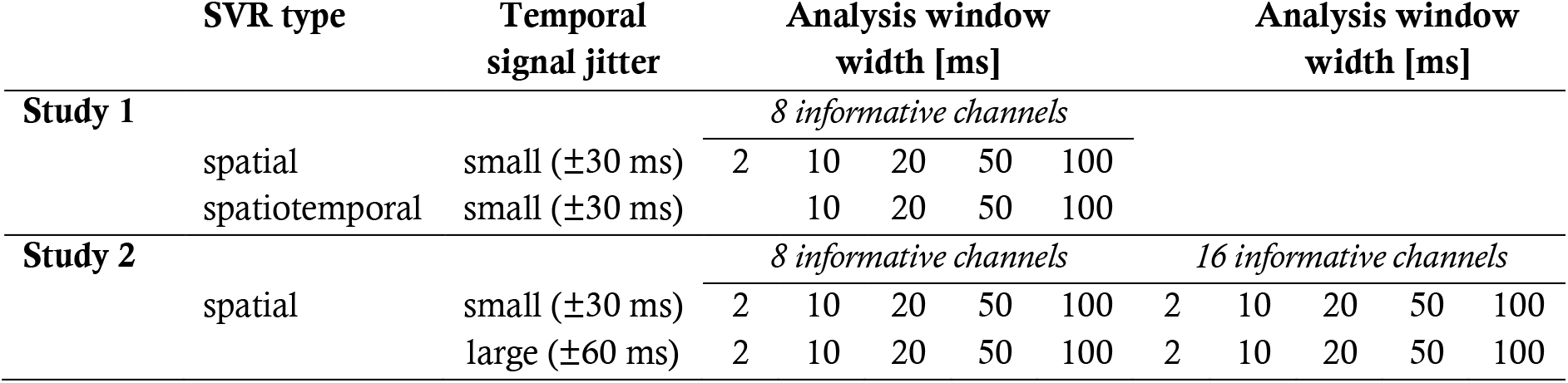
Data generation and decoding analysis settings used for simulation studies 1 and 2.

To simplify the parameter space, we always used the step size of 10 ms with which we moved the analysis time window through the epoch (note that the 2 ms analysis windows required a 2 ms step size to avoid gaps in the resulting information time-course).

We ran the standard ten iterations of a ten-fold cross-validation. For statistical testing, we applied a cluster-based permutation test at *p* < .05 based on the cluster mass statistic (5,000 permutation iterations, cluster-forming alpha = 0.05), as also used in Schubert et al. (2021). The group results represent above-chance Fisher-Z-transformed correlations between predicted label (i.e. the predicted expression of the variable-of-interest) and the real label (i.e. the real expression of the variable-of-interest); or in other words, the decoding performance.

### 2.1 Results

The results showed that all spatial SVR analyses using all analysis time window widths could be used to successfully identify the time period over which the signal was present (Figure 2). The same was true for all spatiotemporal analyses using all time window widths. The results showed that there was no meaningful advantage for using a spatiotemporal over a spatial SVR analysis. The variable-of-interest could be decoded using any analysis time window, including the shortest window, which contained only one data-point per channel per analysis window. The average decoding performance did not differ between analysis approaches (note that the absolute Fisher-transformed Z-value here is somewhat arbitrary and depends on the signal-to-noise ratio (SNR) of the simulated data). However, for both spatial and spatiotemporal SVR, there was some temporal smearing, meaning that time windows earlier in the trial became significant with increasing analysis windows width (more so for the spatiotemporal SVR, where the early informative time points were located as preceding 300 ms). Importantly, this does not reflect higher sensitivity to information, but an artificial projection of information to these early windows. This is due to the way that analysis time windows are constructed: DDTBOX defines a window by the earliest time point included, and it moves the window through the epoch from the start of the epoch to the end of the epoch. As a consequence, any analysis time window that is moved through the epoch becomes significant when predictive information is included at the tail-end of the window. This means, for longer windows, which do not differentiate between information within the window, the true information is, in reality, located at the back (i.e. later time points) of the window, but attributed to the entire window. This could in principle be controlled for by analysing the detailed feature weight structure (including both channels and time points as features); however, as shown here, it can be entirely avoided by using shorter analysis windows.

**Figure 2:**
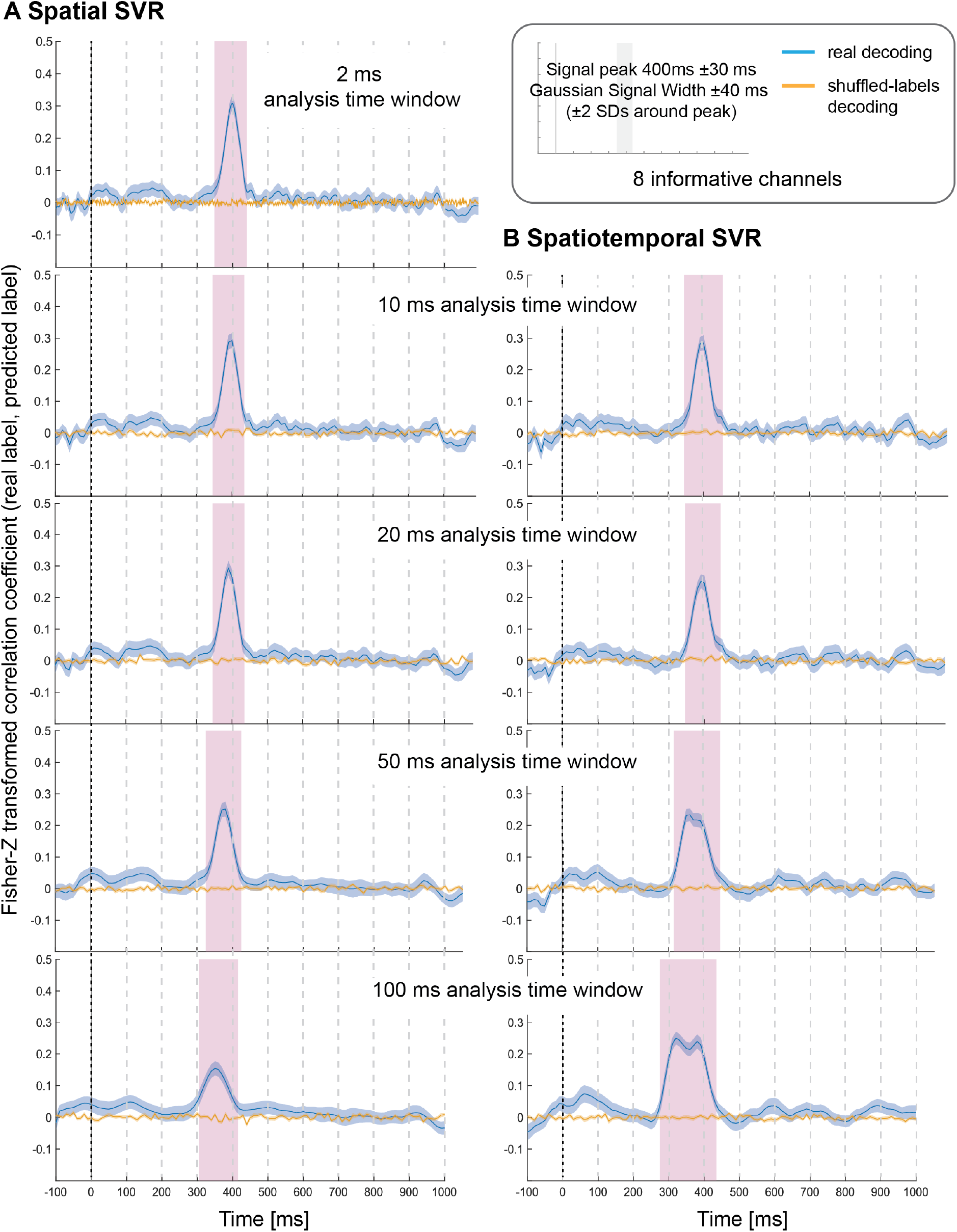
Decoding performance in Simulation Study 1 when using spatial SVR (left panels) and spatiotemporal SVR (right panels), with window widths of 2, 10, 20, 50 and 100 ms. Blue lines denote decoding performance using the original data, orange lines denote decoding performance using permuted data. Shaded regions denote standard errors of the mean (SEMs). Magenta shaded regions denote time windows at which statistically significant above-chance decoding accuracy was found.

Given how similar the results were for spatial and spatiotemporal analyses for this ‘ideal’ dataset (which was characterised by a simple encoding structure of enhanced signal at all informative channels), we opted to only use spatial SVR for the next simulation study. In real data, the precise spatial and temporal location of signal is usually unknown. The second simulation study therefore systematically tested the impact of a) including smaller and larger numbers of information-carrying channels, and b) the presence of smaller or larger temporal jitter in information distribution across these channels.

## 3. Simulation Study 2: Number of informative channels and temporal variance

### 3.1 Methods

#### 3.1.1 Data

Datasets for each experimental condition were generated in the same way as for Simulation Study 1, and we again used N = 37 datasets with the same properties. In this study, we tested four conditions, which incorporated the possible combination of two experimental factors: 1) the number of channels that contained signal (low: 8 channels; high: 16 channels) and 2) the jitter with which the signal was distributed across time in these channels (small: 15 data points / ±30 ms; large: 30 data points / ±60 ms using boxcar distributions as in Simulation Study 1). Table 1 shows all combinations of experimental conditions (note that the condition [16 channels + jitter = ±30 ms] corresponded to the dataset generated for Simulation Study 1). We again applied linear SVR in DDTBOX to analyse all experimental conditions separately.

#### 3.1.1 SVR Analysis

The analysis pipeline and parameters were identical to Simulation Study 1. The only difference was that we only used *spatial* SVR, again in combination with analysis time window widths of 2 ms, 10 ms, 20 ms, 50 ms, and 100 ms (with a step size of 2 ms for the smallest window width, and 10 ms step size for all other window widths). Group-level statistical testing was again conducted using cluster-based permutation tests at *p* < .05 (5,000 permutation iterations, cluster-forming alpha = 0.05) to control for multiple comparisons.

### 3.1 Results

The results again demonstrate that it was possible to decode the variable-of-interest for all experimental conditions with all analysis window widths using spatial SVR (Figure 3 shows analysis window widths of 10 ms and 100 ms; see Supplementary Figures 1-3 for analysis window width 2 ms, 20 ms, and 50 ms). The accuracy and temporal spread of significant decoding results was again highly comparable for the shorter analysis windows (2 ms, 10 ms, 20 ms), which all accurately recovered the underlying time-course of the informative signal. We observed the same temporal smearing of information to earlier time points when using longer analysis time windows, in particular for 100 ms analysis windows, as in the first simulation study.

**Figure 3:**
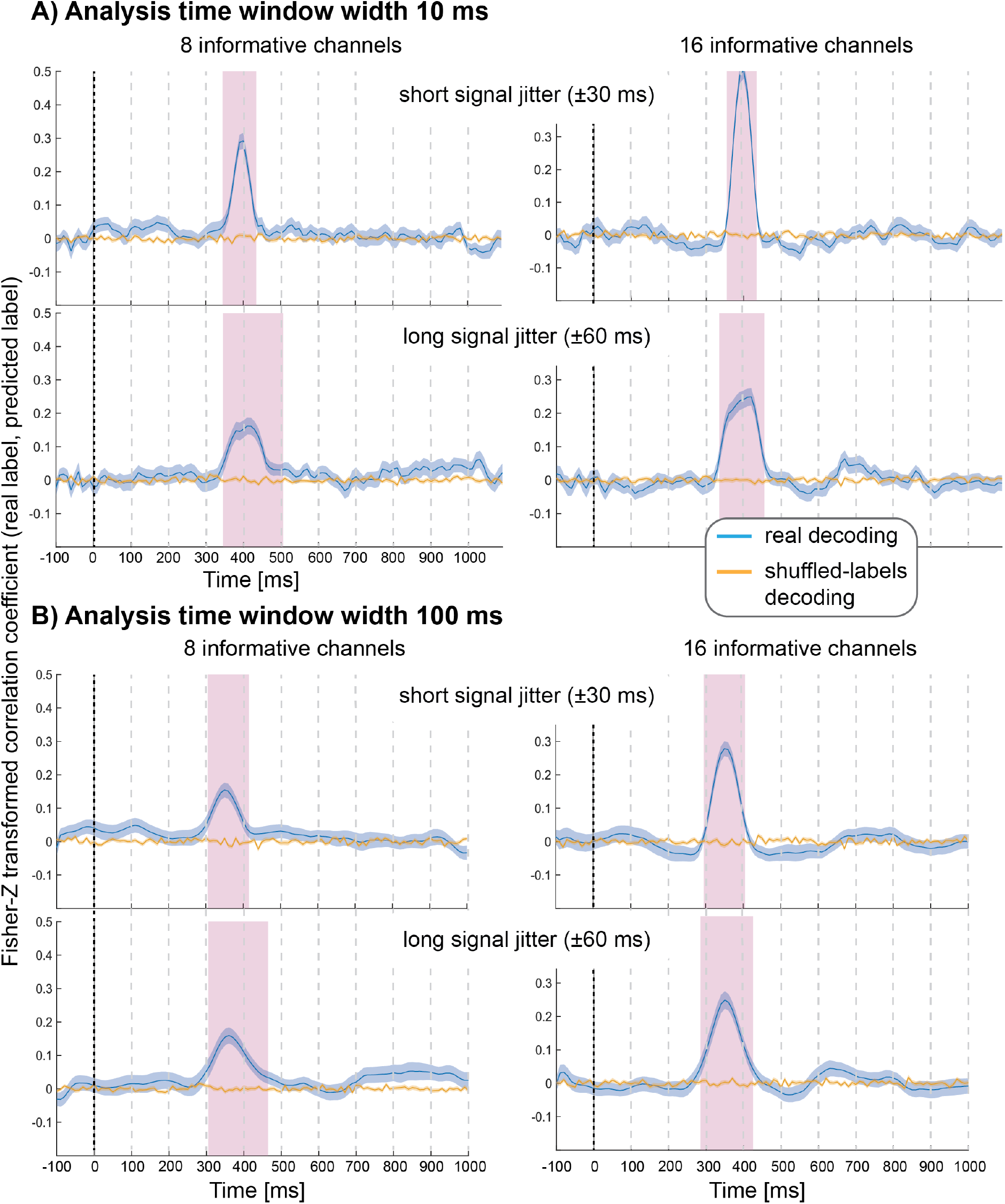
Decoding performance in Simulation Study 2 when using spatial SVR, with window widths of 10 ms (top panels) and 100 ms (bottom panels). Blue lines denote decoding performance using the original data, orange lines denote decoding performance using permuted data. Shaded regions denote standard errors of the mean (SEMs).

As expected, with fewer channels being informative, there was a clear reduction in decoding accuracy; however, it was still possible to significantly predict the variable-of-interest above chance. Introducing a temporal jitter in information distribution for these channels additionally reduced the decoding performance and led to the expected temporal smearing of decoding results across time. Notably, the jitter nevertheless did not prevent the detection of information for any analysis time window.

Magenta shaded regions denote time windows at which statistically significant above-chance decoding accuracy was found.

## 4. Food attribute ERP decoding experiment

Next, we reanalysed a previously published experiment in which participants gave explicit ratings of how tasty and how healthy they perceived visually presented food items to be, while 64-channel EEG was recorded (Schubert et al., 2021). From the analyses reported in the original paper (Schubert et al., 2021), we already learned that decoding was successful using spatiotemporal SVR with 20 ms analysis windows. Here, we reanalysed the data with the same parameters as in Simulation Study 1, i.e. we used both spatial and spatiotemporal linear SVR analyses in combination with 2 ms, 10 ms, 20 ms, 50 ms, and 100 ms analysis time windows. This served to qualitatively compare the simulation results with results from a real EEG study in which the properties of the signal (e.g., the informative channels and the extent to which the signal was jittered) were unknown.

### 4.1 Methods

#### 4.1.1 Participants

Thirty-nine participants were recruited, all right-handed, fluent in written and spoken English, having normal or corrected-to-normal vision, and having no special dietary restrictions or history of eating or feeding disorders. Two participants’ data were excluded because of excessively noisy data. No further exclusions were made based on any other factors. The final sample consisted of 37 participants, ranging from 18 to 36 years old (M = 24.08 years, SD = 4.74; 29 females, 8 males). Participants gave written informed consent before the task and were debriefed and compensated with a voucher valued AUD$20 afterwards. The experiment was approved by the Human Research Ethics Committee (ID1955772) of the University of Melbourne and conducted in accordance with the Declaration of Helsinki.

#### 4.1.2 Stimuli

Stimuli consisted of 174 images of food items from the Food-Pics database (Blechert et al., 2019). Stimuli were selected based on normative palatability ratings and food categorisations from omnivorous participants (Blechert et al., 2019), with the goal of representing a variety of food groups (fruit, vegetables, chocolate, fish, meat, nuts, snacks/meals – sweet and savoury) and a wide range of perceived tastiness. For the full image selection procedure, and questionnaires, and a more detailed description of the procedures, see Schubert et al. (2021).

#### 4.1.3 Task Structure

In the main task, each trial began with a fixation cross (1.5 s), followed by a food image, which was displayed for 2 s (Figure 1 in Schubert et al., 2021). After that, the food image remained on the screen, and in addition a question was displayed below the image – either “How much do you enjoy the taste of this food?” or “How healthy do you consider this food to be?”. Participants answered this question using a continuous sliding scale from “Not at all” to “Very much”, with an underlying range (not visible to participants) of 0 to 100. The end points of the scale were randomly reversed (50% left or right side for each end point) to prevent any motor preparation in the image presentation phase. Participants were given 10 seconds to answer by moving the slider to the position that they believed appropriate and clicking the mouse. If no click was made after 10 seconds, the position of the slider at that time was recorded as a response (which was rarely the case; see Schubert et al., 2021). Each food image was shown twice throughout the experiment, once paired with each question. There were twelve experimental blocks, and in each block only one of the two questions was asked, alternating between blocks. Whether the first block was a taste-rating or health-rating block was randomised across participants. Taken together, the task comprised 348 trials, in which each of the 174 images was rated once for perceived healthiness and once for perceived tastiness. In total, the task took approximately one hour, including EEG setup and pack-up.

#### 4.1.4 EEG Recording and Preprocessing

Electrophysiological activity was recorded using a BioSemi Active II system, with 64 channels, a sample rate of 512 Hz, and recording bandwidth DC-102 Hz. 64 Ag/AgC1 electrodes were attached to a fabric cap according to the International 10-20 system, with four additional electrodes beside and below the left eye (recording the horizontal and vertical electrooculogram), and above the left and right mastoids. Electrode offsets were kept within ±50 μV. Using EEGLab v14.1.2 (Delorme & Makeig, 2004), the data were firstly re-referenced to the average of the left and right mastoids, then high-pass (0.1 Hz) and low-pass (30 Hz) filtered (EEGLab FIR Filter New, default settings). They were segmented into epochs beginning 100 ms before an image was presented and ending 1,000 ms after to capture the first period of visual and semantic processing while avoiding any time periods that already contained activity related to motor preparation. Epochs containing muscle and skin potential artefacts were identified via visual inspection and removed. Excessively noisy channels were removed and interpolated using spherical spline interpolation. An independent components analysis (ICA) was used to identify and remove eye movements, saccades, and blinks. Epochs containing amplitudes at any channel that exceeded ±150 μV were excluded from analyses.

#### 4.1.5 Linear SVR

The linear SVR approach followed exactly the same approach as outlined above, in separate analyses for perceived taste and health. For each analysis, a multivariate regression model was estimated using DDTBOX interfacing LIBSVM (default settings: epsilon-insensitive SVR algorithm; cost parameter C = 0.1) to predict the ratings from the neural data. The average results from all ten iterations of the ten-fold cross-validation procedure were Fisher-Z transformed correlation coefficients for the correlation between the ratings (i.e. labels) and the predicted labels.

As in Simulation Study 1, this analysis was conducted separately using a) spatial SVR and b) spatiotemporal SVR. We again used 2 ms, 10 ms, 20 ms, 50 ms, and 100 ms analysis time windows, moved in steps of 10 ms through the trial (again, only for the 2 ms analysis time window, 2 ms steps were used).

Individual empirical chance results were again created for each participant, obtained by using a shuffled-labels analysis with an identical number of iterations of the same cross-validation procedure with exactly the same data and the same labels (see above). Results from each analysis time window were then tested at group level for statistical significance against the distribution of the empirical chance results by using paired-samples t-tests, corrected for multiple comparisons using cluster-based permutation tests.

### 4.2. Results

The behavioural results and questionnaire results are reported in the original study (Schubert et al., 2021) and are not relevant here.

#### 4.2.1 Taste ratings decoding

Using spatial SVR, taste ratings could be predicted significantly above chance for all analysis time window widths (Figure 4, left side). The general shape of the information time-course was also very similar across all analysis time windows, which confirms the robustness of the approach with real data. The information time-courses for 2 ms and 10 ms windows looked very similar, and the time-course was again smoothed with larger window widths of 20-100 ms because the windows were now moved in overlapping steps. This also meant that the separate significant clusters merged into larger clusters with increasing window width, which is expected since consecutive analyses sampled from overlapping sections of data at larger window width. In general, it can be seen in the figure that the exact significant clusters varied slightly between analysis window width settings. The onset was again systematically propagated forwards in time with larger window width, starting at 540 ms for 10 ms analysis time windows (536 ms for 2 ms window width) and at 470 ms for the 100 ms analysis time windows. For the 100 ms window width, this window included data from between 470 to 570 ms, which means that it is again likely that the tail end of this window started to contain sufficient signal for the analysis to become significant.

**Figure 4:**
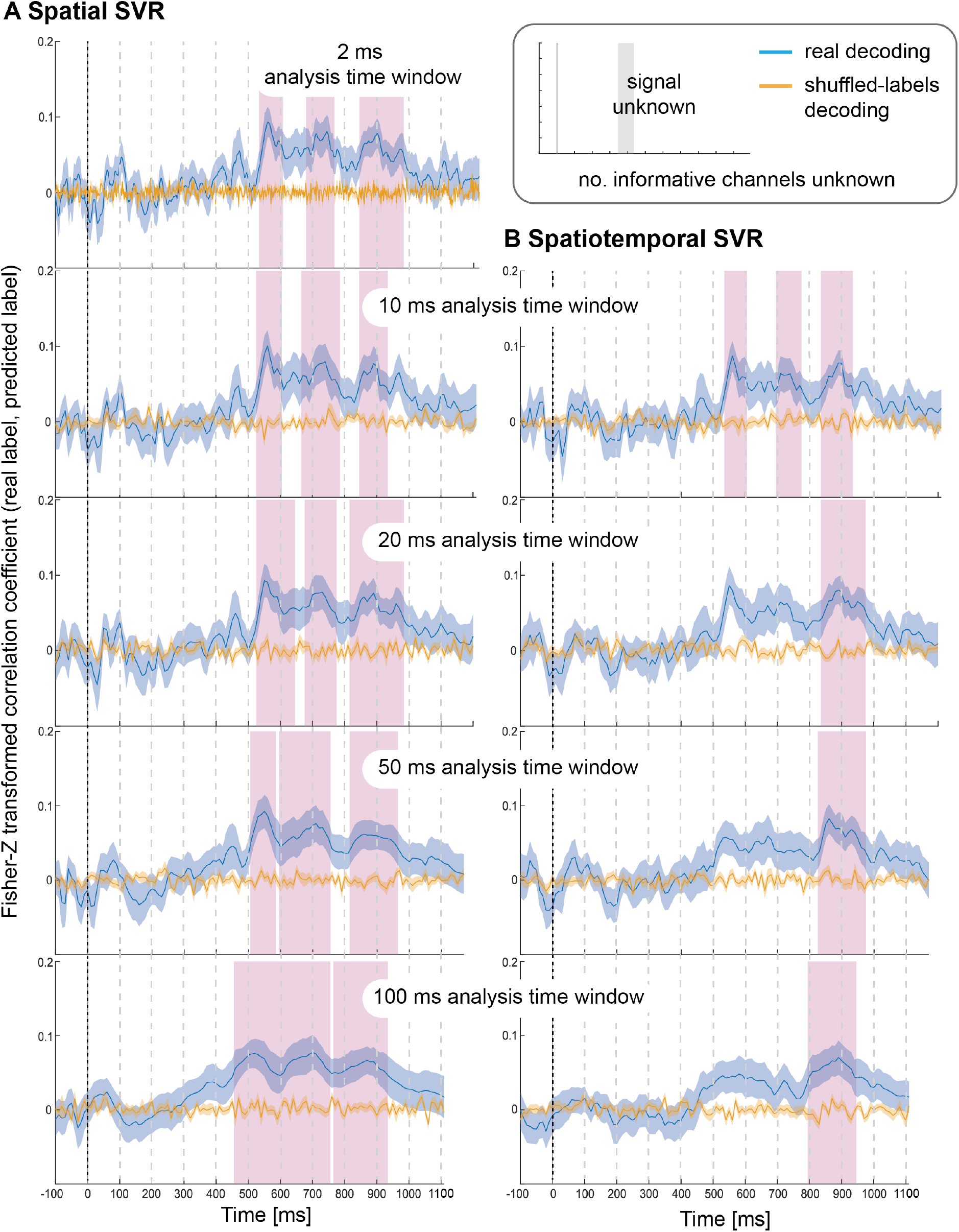
Results for SVR Taste rating decoding. Displayed on the left are results from the spatial decoding analysis, and on the right are the results from the spatiotemporal decoding analysis. Analysis time window widths (from top to bottom): 2 ms (spatial decoding only), 10 ms, 20 ms, 50 ms, 100 ms. Windows were moved in 10 ms steps for all analyses (except for the 2 ms analysis time windows where windows were moved in 2 ms steps). Blue lines are real decoding results (expressed as Fisher-Z-transformed correlation coefficients for the predicted label and the real label, i.e. rating), and yellow lines are chance results from the shuffled-label analyses. Shaded areas represent standard errors of the mean. The food item onset on the screen was at 0 ms (grey line). Significant results after using cluster-corrections for multiple comparisons are highlighted in magenta.

When using spatiotemporal SVR, taste ratings could again be predicted significantly above chance for all analysis time window widths (Figure 4, right side). The general shape of the information time-course for all spatiotemporal analysis time windows was highly similar to their spatial SVR counterparts, and this similarity was strongest for small analysis time windows, in particular at 10 ms (where the onset of the information time-course was 550 ms). Interestingly, some of the earlier clusters only became significant when a 10 ms analysis time window was used, but missed the strict significance threshold for larger analysis time windows. As observed for the simulated data, there was no advantage of using spatiotemporal over spatial analyses. Regardless of window width, the results from the spatiotemporal SVR were slightly lower and more variable across participant datasets than for the spatial SVR, as reflected in fewer and smaller significant clusters in comparison.

#### 4.2.2. Health ratings decoding

Using spatial SVR, the Health ratings could also be predicted significantly above chance for all analysis time windows (Figure 5, left side), although the overall information time-course was noisier than for Taste ratings (which might be due to the use of the rating scale). The general shape of the information time-course was again similar between all analysis time windows, and as before became smoother for larger window widths, with the initial two separate significant clusters merging into one for both 50 ms and 100 ms analysis windows. It can again be seen that whether or not a cluster became significant (after strict cluster-based correction was applied) depended on the analysis time window width used. The same effect of forward-propagating the onset of informative clusters was seen for larger window widths with an onset of 640 ms for 10 ms analysis windows (638 ms for 2 ms) compared to 590 ms for 100 ms analysis time windows (where the first informative window included data from 590 ms to 690 ms).

**Figure 5:**
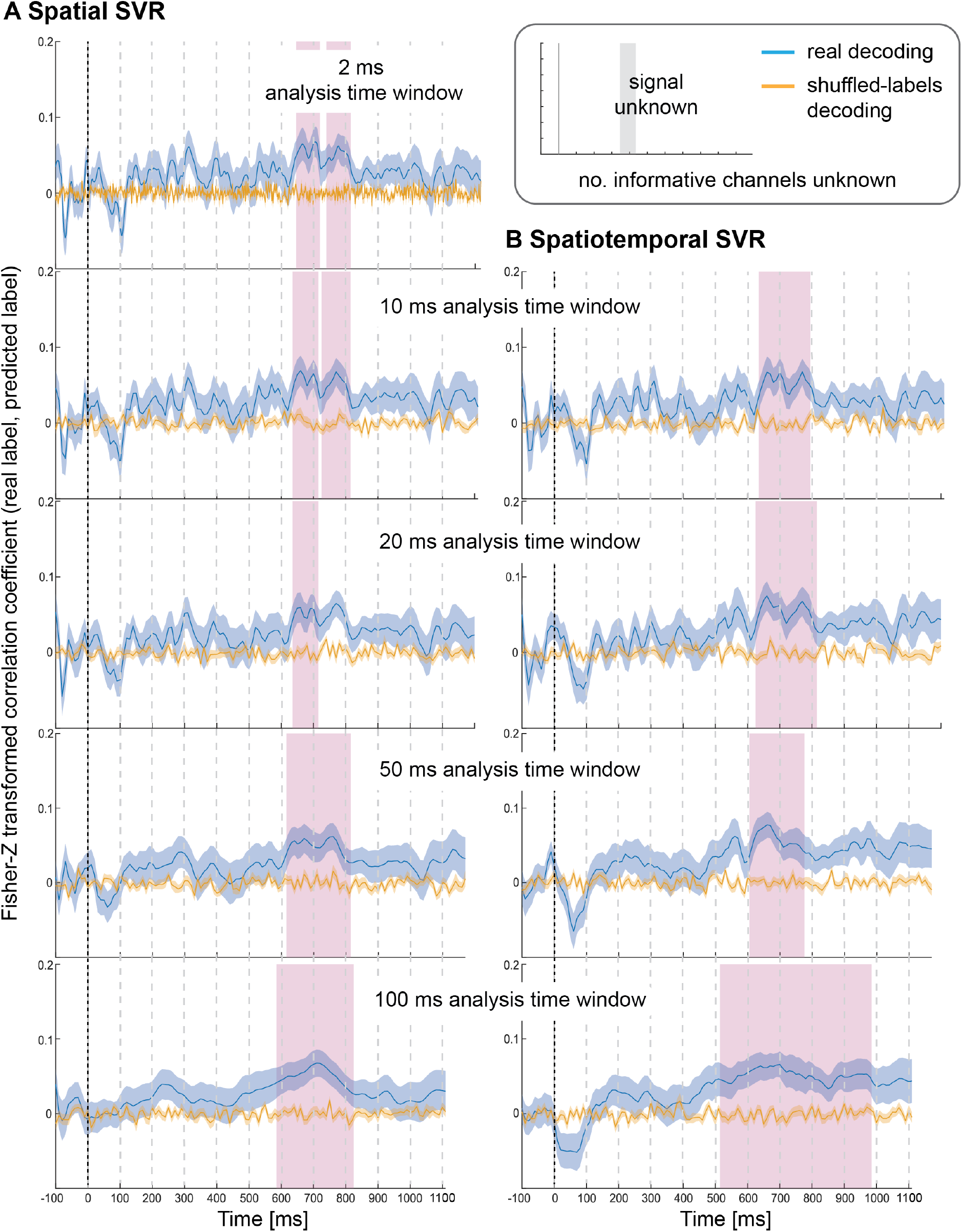
Results for SVR Health rating decoding. Displayed on the left are results from the spatial decoding analysis, and on the right are the results from the spatiotemporal decoding analysis. Analysis time window widths (from top to bottom): 2 ms (spatial decoding only), 10 ms, 20 ms, 50 ms, 100 ms. Windows were moved in 10 ms steps for all analyses (except for the 2 ms analysis time windows where windows were moved in 2 ms steps). Blue lines are real decoding results (expressed as Fisher-Z-transformed correlation coefficients for the predicted label and the real label, i.e. rating), and yellow lines are chance results from the shuffled-label analyses. Shaded areas represent standard error of the mean. The food item onset on the screen was at 0 ms (grey line). Significant results after using cluster-corrections for multiple comparisons are highlighted in magenta.

For the spatiotemporal SVR, there were again no substantial differences compared to the information time-courses from the spatial SVR, in particular for the short 10 ms window width (Figure 5, right side). With 20 ms analysis window width, a larger significant cluster was revealed for spatiotemporal SVR compared to spatial SVR, while with 50 ms windows, it was the other way around, and a slightly larger significant cluster emerged for spatial SVR compared to the spatiotemporal SVR. With 100 ms analysis windows, the significant cluster was substantially larger for spatiotemporal SVR compared to spatial SVR. The shift in onsets of informative clusters again mirrored closely the results from the spatial analysis (however not identical), showing the onset at 640 ms for 10 ms analysis windows, moving to 520 ms for 100 ms analysis time windows (containing data from 520 ms to 620 ms).

In summary, these results show that the findings for these two stimulus dimensions were in general very similar. However, with respect to specific time steps, in some cases a spatial SVR was slightly better, while in other cases, a spatiotemporal SVR was slightly superior – and these subtle differences were again slightly different depending on the analysis time window width.

## 5. Discussion

Support Vector Regression (SVR) is a powerful but underutilised multivariate analysis approach for M/EEG data that still lacks a systematic validation and comprehensive description to become more accessible to the research community. Whereas classification techniques are restricted to predicting categorical variables (i.e. classes), SVR as implemented in DDTBOX (Bode et al., 2019) can predict information about continuous variables from EEG data. Here we used both real and simulated EEG data to demonstrate its usefulness, and we discuss the results of these studies as a guide for future users to help with making decisions on how to tailor SVR in DDTBOX for their specific research questions. We note again that we do not attempt to compare SVR to other classification methods, and a discussion on which specific method might be better suited for specific research questions is also beyond the scope of our paper. Our intention was rather to provide an accessible introduction to SVR in DDTBOX, accompanied with a demonstration of the effects of the most important parameter settings.

The simulation studies allowed us to control the presence of information in the data and showed that both spatiotemporal and spatial analyses generally worked equally well. All analysis time windows, ranging from 2 ms (1 data point per channel) to 100 ms (50 data points per channel), yielded accurate results. When we varied the number of informative channels and the temporal jitter of information for these channels in Simulation Study 2, we observed drops in decoding performance for lower numbers of channels and higher temporal jitter, as expected. Overall, however, we could still recover the information in each condition. We initially chose the parameters (and number of data sets) for the simulation studies deliberately in a way that, in effect, the resulting data was somewhat comparable (although only qualitatively, not quantitatively) to real data.

We then used the EEG data from a recent study by Schubert and colleagues (2021) in which participants rated food attributes to show that the patterns of SVR results were highly similar when predicting continuous taste and health ratings for which the exact distribution of signal in time and space was unknown. This study also showed high similarity between spatial and spatiotemporal SVR (albeit not as similar as in the simulation studies.

### 5.1 Spatial as compared to spatiotemporal SVR

Simulation Study 1 showed that there was no advantage in analysing the full spatiotemporal patterns over spatial patterns for which an average over the data points within each analysis time window was used. This was true for all analysis time window widths, including the long 100 ms window. One explanation for this is that there was no additional information contained in the temporal patterns within windows that could be exploited by spatiotemporal SVR, as we simply added signal to each informative channel across the entire time-period. This might be different in some real data sets, which could potentially have a more complex set of correlations between EEG amplitudes and SVR labels that might vary over time. However, in our real data, we also did not see a clear advantage of spatiotemporal over spatial SVR. This suggests that the temporal information within an analysis time window might only be well-exploited by spatiotemporal methods under specific circumstances, for example, if fast, systematic amplitude fluctuations occur within an analysis window that are not captured by time window-averaged signals.

It should be noted, however, that there might be other real EEG or MEG data sets for which a more fine-grained spatiotemporal approach might be better, but it is difficult to know beforehand if this is the case or not. Even in our real data, it seems that spatial SVR was slightly superior for Taste rating decoding, while spatiotemporal SVR was slightly superior for Health rating decoding, and these differences were additionally slightly more or less pronounced depending on the analysis window width. At this point, it is unclear why such small differences emerge. We therefore believe that it might be premature to only recommend the use of spatial SVR, in particular since previous studies obtained very good results with a spatiotemporal approach (e.g., Bode et al., 2014; Schubert et al., 2020; 2021; Siswandari et al., 2019). On the other hand, there are some disadvantages of using spatiotemporal SVR that should also be taken into account. Firstly, spatiotemporal SVR is computationally more demanding and might require far longer computation time, given the larger number of features that constitute a pattern. Depending on available hardware, there could be processing time advantages in choosing spatial SVR (as we did in Simulation Study 2). Secondly, using spatiotemporal patterns also increases the feature space and, as a side-effect, the risk of overfitting. In particular, if rather large analysis time windows are used (e.g., in our studies, 64 channels in combination with 100 ms analysis window width equals 3,200 features), this can result in an unfavourable ratio between a low number of exemplars (here: epochs) and a high number of features.

### 5.2 Effects of changing the analysis window width

While all analysis windows widths applied here led to significant results, it was obvious that the choice of window width can impact the exact outcomes. Reassuringly, standard analysis window widths between 2 ms and 20 ms led to near-identical results. However, in the real data, some clusters only reached statistical significance (after cluster-based correction) for some analysis window widths. This is likely due to some analysis time windows not quite crossing the p<.05 cluster-forming threshold, rather than substantial differences in decoding performance across window widths. One important implication of this is that whether one obtains significant results for the same data can change by choosing a slightly different analysis window width. Instead of running analyses with all possible window widths, it might therefore be advisable to report the results of the uncorrected significance tests alongside the corrected ones. Another factor influencing the number of statistical tests performed is the length of the chosen epoch. This is also often quite arbitrary and determined by the researchers (ideally informed by the research question). For some questions a long epoch is appropriate, while for others a short epoch is more appropriate. This illustrates that different choices can lead to quite different outcomes – not for the information time-course itself, but for the statistical results. Therefore, it is advised to commit to using an appropriate analysis window width prior to running analyses, and to explain the rationale for using this particular window width.

Another consistent result in our studies was that larger window widths, such as the 100 ms windows, significant effects were observed earlier in the epochs, leading to an over-estimation of how early information was available. This is because analysis time windows in DDTBOX are defined as the time point that comprises the start of the window, meaning that the time period used for analysis extends forward from this time point. Consequently, if there is decodable information at the tail-end of an analysis window (e.g., spanning 390-400 ms when using an analysis window that spans 300-400 ms) then this may still lead to above-chance decoding performance. One way to avoid this is to redefine the analysis window time points as the centre of the window, but even then, some unavoidable temporal smearing would remain. The better solution might be to avoid such large analysis windows altogether; this also makes sense as the advantage of the excellent temporal resolution of EEG is lost when such long windows are used. The same basic problem emerges, of course, when the data is substantially down-sampled first (which is basically the same as using a spatial SVR with large analysis window width).

Interestingly, in our studies, it was possible to use analysis time windows as small as 2 ms (containing only one data point from each channel), or 10 ms (containing 5 data points) without an obvious drop in decoding performance. This is encouraging as it implies that one can indeed make use of the excellent temporal resolution of EEG and derive quite precise information time-courses. However, it should be kept in mind that in real data, there might be differences in performance between time analysis windows of different widths depending on the underlying cognitive process (and the duration of a stable neural pattern). Long duration analysis time windows might be ill-suited to capture fast, short-duration neural processes, because each analysis time window would include a short period of signal intermixed with a larger period containing only noise. Whether the reverse is also true, and short analysis time windows may have problems capturing longer and sustained processes, is not so clear. This might depend on the encoding of the cognitive process in the neural pattern. Our results suggest that shorter analysis time windows had no problem detecting signals that were variable in their timing, both in the real data as well as in Simulation Study 2 where a larger extent of temporal jitter was used. In fact, the use of temporal generalisation matrices (TGMs) in classification studies relies on the idea that a relatively short analysis windows can capture subsequent aspects of the neural signature of the same sustained cognitive process. This approach should further be sensitive to characterise changes in these informative patterns over time, which can inform the researchers that the nature of a representation has changed over time (e.g., King & Dehaene, 2014; Blom et al., 2020; 2021). This logic should also translate to SVR approaches.

However, very little is known about the nature of more abstract neural representations of cognitive processes, potentially encoded in prefrontal and parietal cortices, compared to sensory-based representations or motor representations. The few existing studies using linear SVR have rarely reported systematic comparisons of analysis time window widths. It therefore remains to be understood whether there are more or less optimal window widths for specific abstract representations, or other sustained cognitive processes, which might not be associated with strong, sustained signals over time (e.g., keeping an item in memory; or activating an emotion regulation goal).

### 5.3 Limitations

It should be noted that this study was designed as a first (and somewhat belated) proof-of-principle for the use of linear SVR as a multivariate approach to analysing ERP data using DDTBOX. As such, it captured the most important parameter settings in DDTBOX, to give users a good idea about their effect, but we did not attempt to provide a complete exploration of the entire parameter space. DDTBOX (through LIBSVM) allows users to deviate from standard SVR settings, for example, to use non-linear kernels, which might be better suited for specific problems. There are also a multitude of other settings, which we did not explore here: we did not cover the entire range of other possible analysis window widths; we did not compare the effects of different cost parameters; and we did not compare different approaches for the cross-validation procedure (nor the number of iterations; nor nested cross-validation to optimise the cost parameter). The main reason is that these settings are less frequently used in the field. We also did not compare different strategies to control for the multiple comparison problem for group-statistical testing, but we provide an overview on our Wiki. We also did not analyse the feature weights here which can give some insights into the contribution of single channels to the prediction (if controlled appropriately, see Haufe et al., 2014) as this goes beyond the scope of this paper. There are also alternative approaches to quantifying the outcome of the regression analyses to the normalised correlations used here (such as normalised mean absolute, or squared, errors) that are not yet implemented in DDTBOX, but users might prefer calculating themselves. However, the approach presented here is most frequently used in the literature and has the advantage that it is readily interpretable.

Regarding the simulated data, we also restricted our simulation studies to two conditions, comparing the number of informative channels and the temporal jitter, respectively, to provide some insight into their effects. Since the data structure and information time-course of real EEG data is usually unknown, there is scope for exploring much more extreme conditions; however, in combination with the other factors in our studies (e.g., SVR types and analysis window widths), this would have led to a very high number of analyses to report in a single paper (and results are less relevant for typical users). Similarly, there are many other, unexplored factors which are potentially of interest for future simulation studies, including: the relationship between sample size, number of exemplars/epochs, and statistical power to detect effects of different sizes; the mapping of continuous variables of interest to single-trial ERP amplitudes, and how different underlying distributions of this variable might impact predictability; and the impact of the SNR on predictive performance. We have further refrained from comparing the SVR approach to other, related multivariate approaches, such as support vector classification, classification using other methods (such as linear discriminant analysis; Carlson, Schrater & He 2003), extensions to these approaches that extract a continuous output variable (such as the distance-to-bound approach, Ritchie & Carlson 2016), and other multivariate regression approaches (e.g., Parra et al., 2005). Such an exploration was beyond the scope of the current paper. However, given the growing diversity of methods in this space, this would indeed be a valuable exercise for the future.

### 5.4 Conclusions

Multivariate classification approaches have become a powerful tool in the toolbox of cognitive neuroscience, but a key constraint of such approaches is that they are often used to predict a categorical output variable (i.e. distinct classes). Classifying between high and low values of a continuous variable using median-splits, however, suffers from reduced statistical power. Here, we present a guide for users on how to set up a SVR analysis as implemented in DDTBOX for the prediction of continuous variables of interest from multivariate patterns of ERPs, as measured by EEG. To help users to better understand the effects of different parameter settings, we conducted a series of analyses. These demonstrate that SVR analysis can predict a continuous variable in both simulated and real EEG data. Results were highly similar for spatial and spatiotemporal SVR and stable across a variety of analysis time window widths, including very short analysis windows containing only one data point per channel. Some temporal smearing occurred with larger analysis time windows. SVR could successfully predict the variables of interest even with a smaller number of informative channels and larger temporal signal jitters. These results, together with previously published demonstrations of successful prediction of a variety of continuous variables, suggest that SVR constitutes a useful data-driven analysis approach, which allows for subtler variations of variables of interest to be predicted from distributed patterns of neural activity at high temporal resolution. We hope that, together with our previous publication on DDTBOX (Bode et al., 2019), this paper will provide users with a valuable resource to optimally conduct SVR analyses in their own data.

## Acknowledgements

HH acknowledges support from the Australian Research Council (DP180102268 and FT200100246). DF was supported by the Australian Research Council Discovery Early Career Researcher Award (ARC DE220101508). We thank Mackenzie Murphy for help with data collection. The funding bodies had no input into the design, results or interpretation of this work.

## References

1. Allefeld, C., Görgen, K., Haynes, J.D., 2016. Valid population inference for information-based imaging: From the second-level t-test to prevalence inference. Neuroimage, 141, 378–392. doi: 10.1016/j.neuroimage.2016.07.040.

2. Bai, O., Lin, P., Vorbach, S., Li, J., Furlani, S., Hallett, M., 2007. Exploration of computational methods for classification of movement intention during human voluntary movement from single trial EEG. Clin. Neurophysiol., 118, 2637– 2655. doi: 10.1016/j.clinph.2007.08.025.

3. Billing, A.J., Davis, M.H., Carlyon, R.P., 2018. Neural decoding of bistable sounds reveals an effect of intention on perceptual organization. J. Neurosci., 38(11), 2844–2853. doi: 10.1523/JNEUROSCI.3022-17.2018.

4. Blechert, J., Lender, A., Polk, S., Busch, N. A., & Ohla, K., 2019. Food-pics_extended—an image database for experimental research on eating and appetite: additional images, normative ratings and an updated review. Front. Psychol., 10, 307. doi: 10.3389/fpsyg.2019.00307.

5. Blom, T., Bode, S., Hogendoorn, H., 2021. The time-course of prediction formation and revision in human visual motion processing. Cortex, 138, 191–202. doi: 10.1016/j.cortex.2021.02.008.

6. Blom, T., Feuerriegel, D., Johnson, P., Bode, S., Hogendoorn, H., 2020. Predictions drive neural representations of visual events ahead of incoming sensory information. Proc. Natl. Acad. Sci. U.S.A., 117(13), 7510–7515. doi: 10.1073/pnas.1917777117.

7. Bode, S., Bennett, D., Stahl, J., Murawski, C., 2014. Distributed patterns of event-related potentials predict subsequent ratings of abstract stimulus attributes. PLoS ONE, 9(10), e109070. doi: 10.1371/journal.pone.0109070.

8. Bode, S., Feuerriegel, D., Bennett, D., Alday, P.M., 2019. The Decision Decoding ToolBOX (DDTBOX)–A multivariate pattern analysis toolbox for event-related potentials. Neuroinform., 17(1), 27–42. doi: 10.1007/s12021-018-9375-z.

9. Bode, S., Sewell, D.K., Lilburn, S., Forte, J., Smith, P.L., Stahl, J., 2012. Predicting perceptual decision biases from early brain activity. J. Neurosci., 32(36), 12488– 12498. doi: 10.1523/JNEUROSCI.1708-12.2012.

10. Bullmore, E.T., Suckling, J., Overmeyer, S., Rabe-Hesketh, S., Taylor, E., Brammer, M. J., 1999. Global, voxel, and cluster tests, by theory and permutation, for a difference between two groups of structural MR images of the brain. IEEE Trans. Med. Imaging, 18(1), 32–42. doi: 10.1109/42.750253.

11. Carlson, T.A., Grootswagers, T., Robinson, A.K., 2020. An introduction to time-resolved decoding analysis for M/EEG. arXiv, 1905.04820.

12. Carlson, T.A., Schrater, P., He, S., 2003. Patterns of activity in the categorical representations of objects. J. Cogn. Neurosci., 15(5), 704–717. doi: 10.1162/089892903322307429.

13. Chang, C.C., Lin, C.J., 2011. LIBSVM: a library for support vector machines. ACM Trans. Intell. Syst. Technol., 2, 1–27. doi: 10.1145/1961189.1961199.

14. Combrisson, E., Jerbi, K., 2015. Exceeding chance level by chance: The caveat of theoretical chance levels in brain signal classification and statistical assessment of decoding accuracy. J. Neurosci. Methods, 250, 126–136. doi: 10.1016/j.jneumeth.2015.01.010.

15. Contini, E.W., Wardle, S.G., Carlson, T.A., 2017. Decoding the time-course of object recognition in the human brain: From visual features to categorical decisions. Neuropsychologia, 105, 165–176. doi: 10.1016/j.neuropsychologia.2017.02.013.

16. Delorme, A., Makeig, S., 2004. EEGLAB: an open source toolbox for analysis of single-trial EEG dynamics including independent component analysis. J. Neurosci. Methods, 134, 9–21. doi: 10.1016/j.jneumeth.2003.10.009.

17. Fahrenfort, J.J., van Driel, J., van Gaal, S., Olivers, C.N.L., 2018. From ERPs to MVPA Using the Amsterdam Decoding and Modeling Toolbox (ADAM). Front. Neurosci., 12, 368. doi: 10.3389/fnins.2018.00368.

18. Fan, R.E., Chang, K.-W., Hsieh, C.J., Wang, X.R., Lin, C.J., 2008. LIBLINEAR: a library for large linear classification. J. Machine Lean. Res., 9, 1871–1874.

19. Friston, K., Holmes, A., Price, C., Buchel, C., Worsley, K., 1999. Multisubject fMRI studies and conjunction analysis. Neuroimage, 10, 385–396. doi: 10.1006/nimg.1999.0484.

20. Gevins, A., 1989. Dynamic functional topography of cognitive tasks. Brain Topogr. 2, 37–56. doi: 10.1007/BF01128842.

21. Gramfort, A., Luessi, M., Larson, E., Engemann, D., Strohmeier, D., Brodbeck, C., Goj, R., Jas, M., Brooks, T., Parkkonen, L., Hämäläinen, M., 2013. MEG and EEG data analysis with MNEPython. Front. Neurosci, 7. doi: 10.3389/fnins.2013.00267.

22. Grootswagers, T., Wardle, S.G., Carlson, T.A., 2016. Decoding dynamic brain patterns from evoked responses: A tutorial on multivariate pattern analysis applied to time series neuroimaging data. J. Cogn. Neurosci., 29, 677–697. doi: 10.1162/jocn_a_01068.

23. Hanke, M., Halchenko, Y.O., Sederberg, P.B., Hanson, S.J., Haxby, J.V., Pollman, S., 2009. PyMVPA: A python toolbox for multivariate pattern analysis of fMRI data. Neuroinform., 7, 37–53. doi: 10.1007/s12021-008-9041-y.

24. Hastie, T., Tibshirani, R., Friedman, J., 2009. The elements of statistical learning: Data mining, inference, and prediction. Heidelberg: Springer Science & Business Media.

25. Haufe, S., Meinecke, F., Görgen, K., Dähne, S., Haynes, J.D., Blankertz, B., Bießmann, F., 2014. On the interpretation of weight vectors of linear models in multivariate neuroimaging. Neuroimage, 87, 96–110. doi: 10.1016/j.neuroimage.2013.10.067.

26. Hebart, M.N., Görgen, K., Haynes, J.D., 2015. The Decoding Toolbox (TDT): A versatile software package for multivariate analyses of functional imaging data. Front. Neuroinform., 8, 88. doi: 10.3389/fninf.2014.00088.

27. Jach, H.K., Feuerriegel, D., Smillie, L.D., 2020. Decoding personality trait measures from resting EEG: An exploratory report. Cortex, 130, 158–171. doi: 10.1016/j.cortex.2020.05.013.

28. King, J.R., Dehaene, S., 2014. Characterizing the dynamics of mental representations: The temporal generalization method. Trends Cogn. Sci., 18, 203–210. doi: 10.1016/j.tics.2014.01.002.

29. Korjus, K., Uusberg, A., Uusberg, H., Kuldkepp, N., Kreegipuu, K., Allik, J., … & Aru, J., 2015. Personality cannot be predicted from the power of resting state EEG. Frontiers in Human Neuroscience, 9, 63. doi: 10.3389/fnhum.2015.00063.

30. Lan, Z., Müller-Putz, G.R., Wang, L., Liu, Y., Sourina, O., Scherer, R., 2016. Using Support Vector Regression to estimate valence level from EEG. *IEEE International Conference on Systems*, Man, and Cybernetics (SMC), 002558–002563. doi: 10.1109/SMC.2016.7844624.

31. Li, J.-W., Wang, Y.-H., Wu, Q., Wei, Y.-F., An, J.-L., 2008. EEG source localization of ERP based on multidimensional support vector regression approach. IEEE International Conference on Machine Learning and Cybernetics, 1238–1243. doi: 10.1109/ICMLC.2008.4620594.

32. MacCallum, R. C., Zhang, S., Preacher, K. J., Rucker, D. D. (2002). On the practice of dichotomization of quantitative variables. Psychol. Methods, 7(1), 19. doi:10.1037/1082-989x.7.1.19

33. Maris, E., Oostenveld, R., 2007. Nonparametric statistical testing of EEG- and MEG-data. J. Neurosci. Methods, 164, 177–190. doi: 10.1016/j.jneumeth.2007.03.024.

34. Meyers, E.M., 2013. The neural decoding toolbox. Front. Neuroinform., 7, 8. doi: 10.3389/fninf.2013.00008

35. Oostenveld, R., Fries, P., Maris, E., Schoffelen, J.-M., 2011. FieldTrip: Open source software for advanced analysis of MEG, EEG, and invasive electrophysiological data. Comput. Intell. Neurosci., 156869. doi: 10.1155/2011/156869.

36. Oosterhof, N.N., Connolly, A.C., Haxby, J.V., 2016. CoSMoMVPA: Multi-modal multivariate pattern analysis of neuroimaging data in Matlab/GNU Octave. Front. Neuroinform., 10, 27. doi: 10.3389/fninf.2016.00027.

37. Parra, L.C., Spence, C.D., Gerson, A.D., Sajda, P., 2005. Recipes for the linear analysis of EEG. Neuroimage, 28(2), 326–341. doi: 10.1016/j.neuroimage.2005.05.032.

38. Perrin, F., Bertrand, O., Pernier, J., 1987. Scalp current density mapping: value and estimation from potential data. IEEE Trans. Biomed. Eng., 34, 283–288. doi: 10.1109/tbme.1987.326089.

39. Pernier, J., Perrin, F., Bertrand, O., 1988. Scalp current density fields: Concepts and properties. Electroencephalogr. Clin. Neurophysiol., 69, 385–389. doi: 10.1016/0013-4694(88)90009-0.

40. Ritchie, J.B., Carlson, T.A., 2016. Neural decoding and “inner” psychophysics: A distance-to-bound approach for linking mind, brain, and behavior. Front. Neurosci., 10, 190. doi: 10.3389/fnins.2016.00190.

41. Sabbagh, D., Ablin, P., Varoquaux, G., Gramfort, A., Engemann, D.A., 2020. Predictive regression modeling with MEG/EEG: from source power to signals and cognitive states. Neuroimage, 222, 116893. doi: 10.1016/j.neuroimage.2020.116893.

42. Sato, J.R., Costafreda, S., Morettin, P.A., Brammer, M.J., 2008. Measuring time series predictability using support vector regression. Commun. Stat. Simul. Comput., 37, 1183–1197. doi: 10.1080/03610910801942422.

43. Schrouff, J., Rosa, M.J., Rondina, J.M., Marquand, A.F., Chu, C., Ashburner, J., et al., 2013. PRoNTo: Pattern recognition for neuroimaging toolbox. Neuroinform., 11, 319–337. doi: 10.1007/s12021-013-9178-1.

44. Schubert, E., Agathos, J.A., Brydevall, M., Feuerriegel, D., Koval, P., Morawetz, C., Bode, S., 2020. Neural patterns during anticipation predict emotion regulation success for reappraisal. Cogn. Affect. Behav. Neurosci., 20(4), 888–900. doi: 10.3758/s13415-020-00808-2.

45. Schubert, E., Rosenblatt, D., Eliby, D., Kashima, Y., Hogendoorn, H., Bode, S., 2021. Decoding explicit and implicit representations of health and taste attributes of foods in the human brain. Neuropsychologia, 162, 108045. doi: 10.1101/2021.05.16.444383v1.

46. Siswandari, Y., Bode, S., Stahl, J., 2019. Performance monitoring beyond choice tasks: The time course of force execution monitoring investigated by event-related potentials and multivariate pattern analysis. Neuroimage, 197, 544–556. doi: 10.1016/j.neuroimage.2019.05.006.

47. Treder, M.S., 2020. MVPA-Light: A classification and regression toolbox for multi-dimensional data. Front. Neurosci., 14, 289. doi: 10.3389/fnins.2020.00289.

48. Trübutschek, D., Marti, S., Ojeda, A., King, J.R., Mi, Y., Tsodyks, M., Dehaene, S., 2017, A theory of working memory without consciousness or sustained activity. eLife, 6, e23871. doi: 10.7554/eLife.23871.

