## Supplementary Material for "Decoding continuous variables from event-related potential (ERP) data with linear support vector regression (SVR) using the Decision Decoding Toolbox (DDTBOX)"

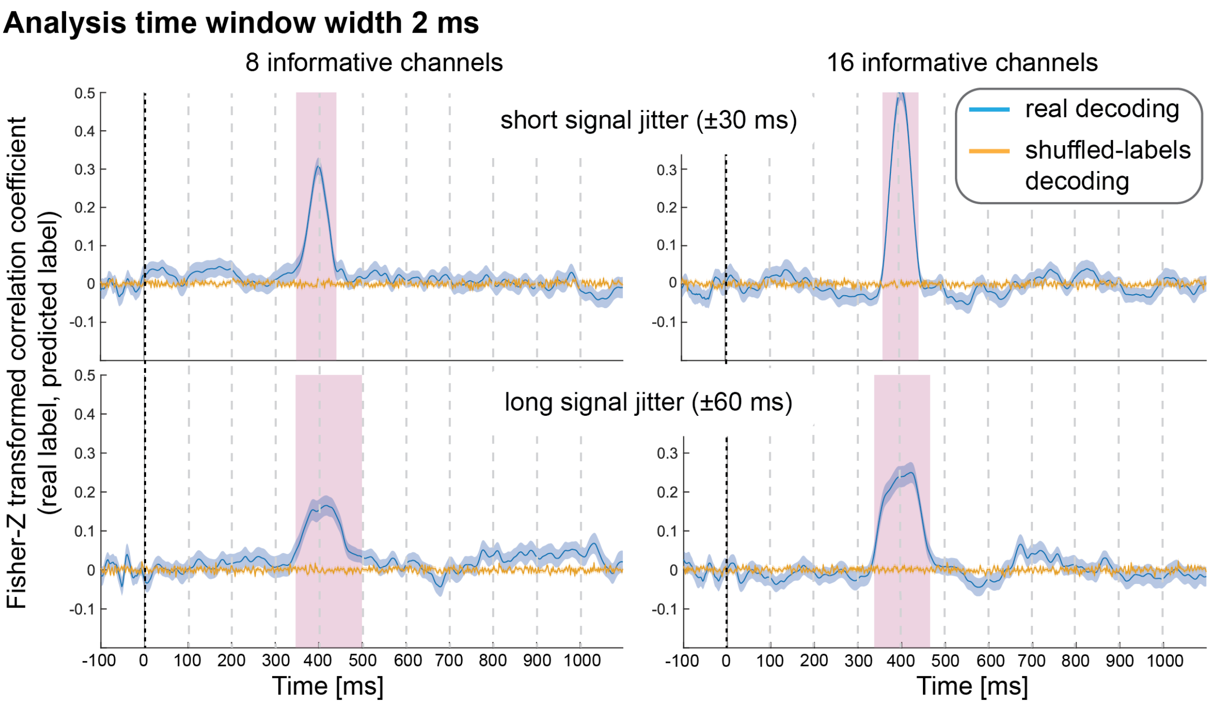


**Supplementary Figure 1**: Decoding performance in Simulation Study 2 when using spatial SVR, with window widths of 2 ms. Blue lines denote decoding performance using the original data, orange lines denote decoding performance using permuted data. Shaded regions denote standard errors of the mean (SEMs). Magenta shaded regions denote time windows at which statistically significant above-chance decoding accuracy was found.

### Analysis time window width 20 ms

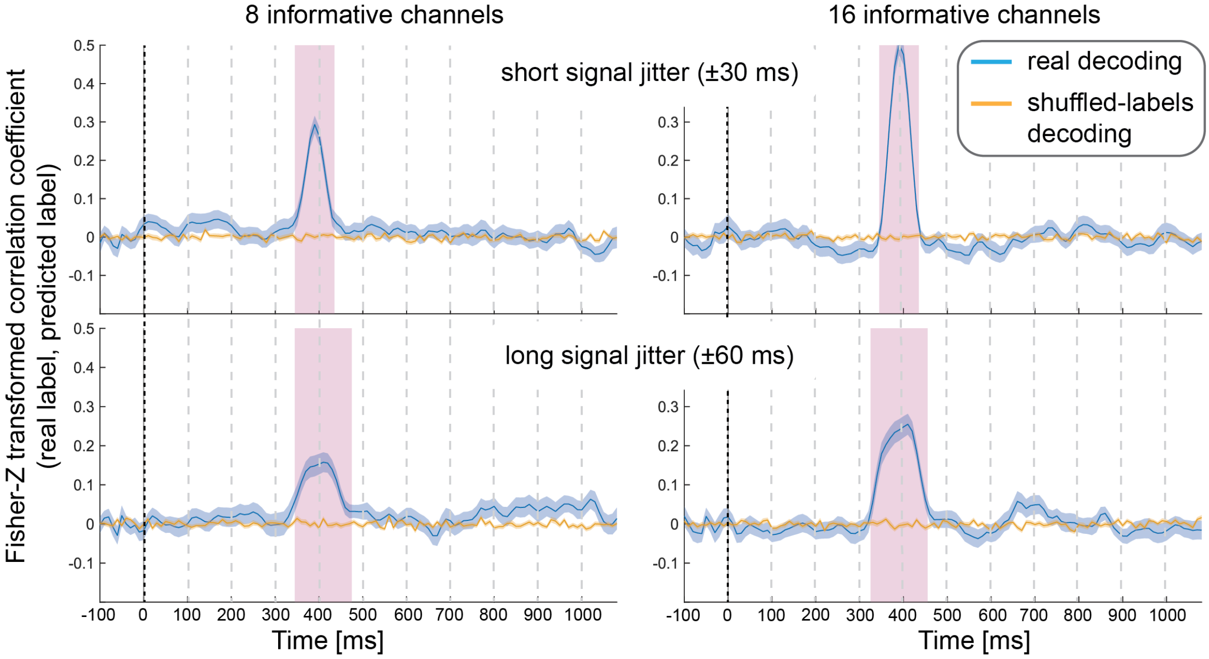


**Supplementary Figure 2**: Decoding performance in Simulation Study 2 when using spatial SVR, with window widths of 20 ms. Blue lines denote decoding performance using the original data, orange lines denote decoding performance using permuted data. Shaded regions denote standard errors of the mean (SEMs). Magenta shaded regions denote time windows at which statistically significant above-chance decoding accuracy was found.

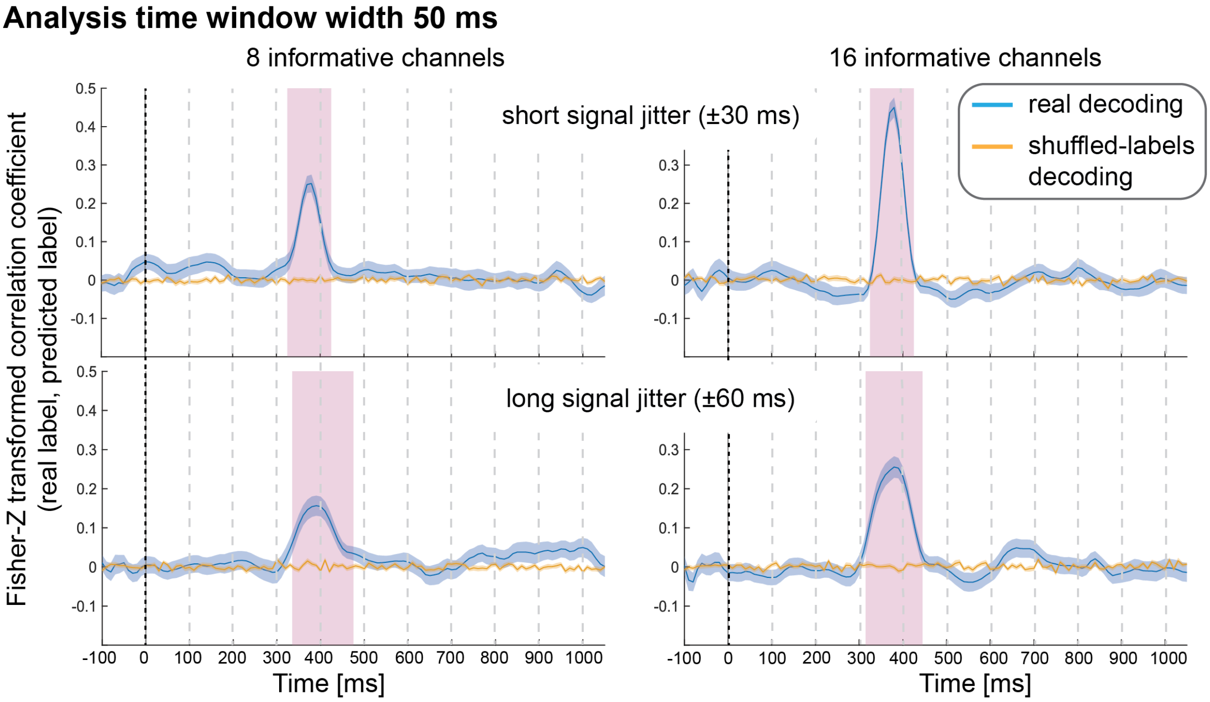


**Supplementary Figure 3**: Decoding performance in Simulation Study 2 when using spatial SVR, with window widths of 50 ms. Blue lines denote decoding performance using the original data, orange lines denote decoding performance using permuted data. Shaded regions denote standard errors of the mean (SEMs). Magenta shaded regions denote time windows at which statistically significant above-chance decoding accuracy was found.
